# Ventricular lymphatic vessel density correlates with mild cardiac remodelling and preserved heart function in moderate murine aortic valve stenosis

**DOI:** 10.1101/2024.07.06.602334

**Authors:** Katrin Becker

**Author notes:** Mailing address: Institute for Cardiovascular Sciences, Medical Faculty and University Hospital Bonn, University Bonn, Venusberg-Campus 1, 53127 Bonn, Germany. c) Author contribution* K. B. conducted conceptualization, investigation, data curation - formal analysis, visualization, writing – original draft, review and editing.

## Abstract

Aortic valve stenosis (AS) is one of the most frequent heart valve diseases in the western world, but aortic valve replacement (AVR) remains the only therapeutic option. While one of the most common complications of AS is heart failure, the early disease phase is asymptomatic with preserved cardiac function. Increased ventricular lymphatic vessel density is part of myocardial remodelling upon cardiac pressure overload and an important adaptation mechanism to maintain heart function after myocardial infarction, while in heart failure patients, circulating lymphatic growth factor levels and ventricular lymphatic vessel numbers are altered. Lymphangiogenesis is enhanced in endocardium of human stenotic aortic valves, however, ventricular lymphatic vessels have not been investigated in AS, yet.

Therefore, aim of this study was to in a well-characterized mouse model of moderate AS analyse the link between density of ventricular lymphatic vessels, cardiac remodelling and function.

## Introduction

Aortic valve stenosis (AS) is one of the most frequent heart valve diseases in the western world, but aortic valve replacement (AVR) remains the only therapeutic option. While one of the most common complications of AS is heart failure, the early disease phase is asymptomatic with preserved cardiac function[1]. Increased ventricular lymphatic vessel density is part of myocardial remodelling upon cardiac pressure overload[2] and an important adaptation mechanism to maintain heart function after myocardial infarction[3], while in heart failure patients, circulating lymphatic growth factor levels[4] and ventricular lymphatic vessel numbers[5] are altered. Lymphangiogenesis is enhanced in endocardium of human stenotic aortic valves[6], however, ventricular lymphatic vessels have not been investigated in AS, yet.

Therefore, aim of this study was to in a well-characterized mouse model of moderate AS[7] analyse the link between density of ventricular lymphatic vessels, cardiac remodelling and function.

## Methods

### In vivo *animal experiments*

Twelve eight-week-old male C57Bl/6J mice (Janvier Labs, France) were randomly assigned to wire injury (WI) to induce moderate AS, or sham surgery, as published previously[7]. Echocardiography was conducted at baseline, two and four weeks after surgery. Inclusion criterion: Development of AS (blood flow increase to >1.5 of baseline or >2000 mm/sec); exclusion criteria: Severe aortic valve regurgitation, or death prior to endpoint (here: one sham, three WI animals). Model limitations are differences between human and murine lymphatic vasculature [2], mouse strains and sexes, wherefore results cannot be generalized.

The animals were sacrificed four weeks after surgery (isoflurane overdose) and immediately perfused with phosphate buffered saline (**figure 1 A**).

**Figure 1.**
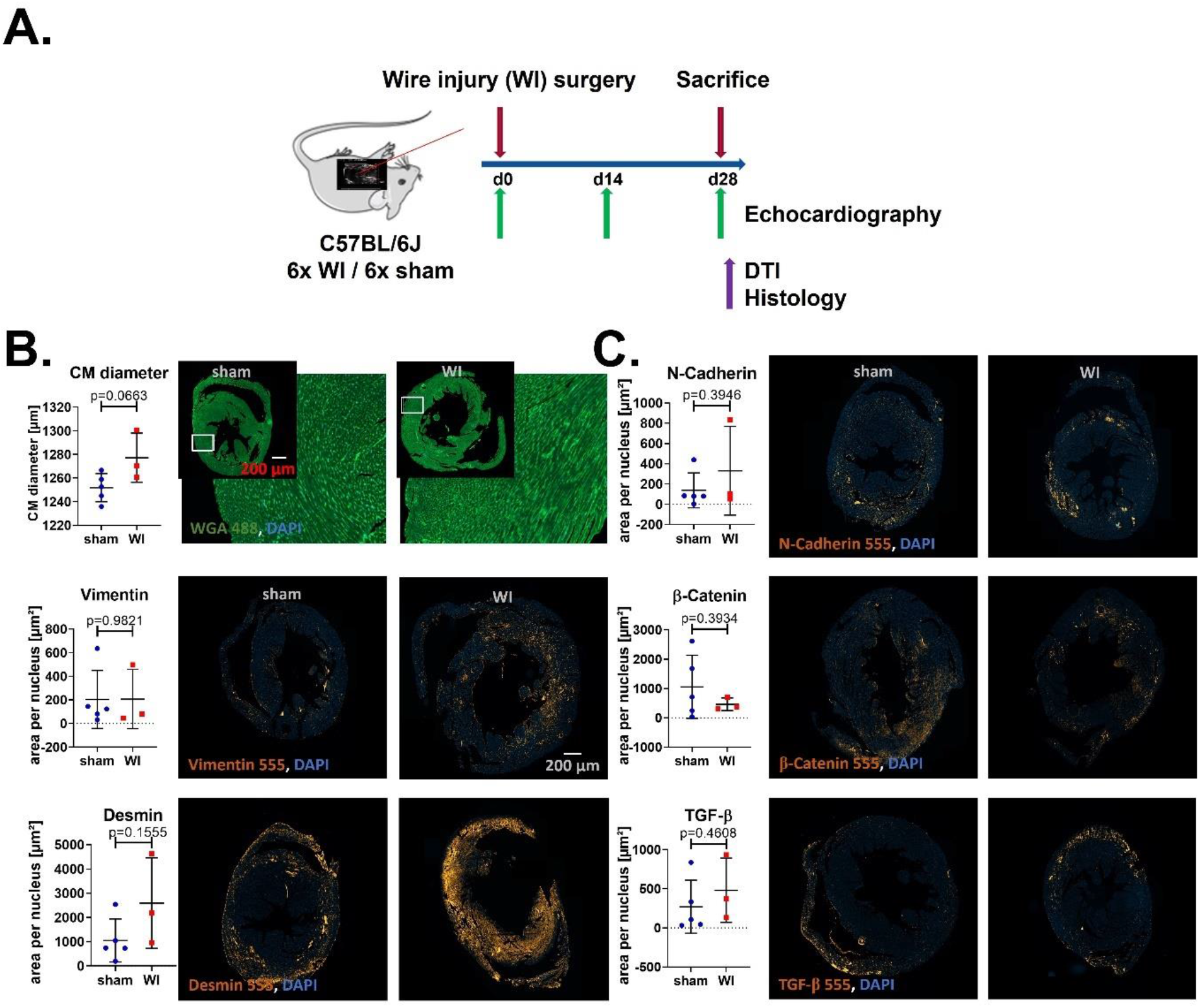
**A)** Wire injury surgery (WI) was performed in 8-week-old male C57BL/6J mice, with echocardiography at baseline, and after 14 and 28 days to verify manifestation of AS. Ex vivo diffusion tensor imaging (DTI) of explanted hearts was done at 11 T. For histology, mid-ventricular 4 μm heart slices were stained as described previously[7]. For statistical analysis, Mann-Whitney test was used. **B + C)** Histologic analysis of cardiac remodeling: Cardiomyocyte (CM) surfaces: Wheat germ agglutinin (WGA). Fibrosis marker: Vimentin; marker of cardiomyocyte remodeling: Desmin (**B**). Markers for signaling pathways underlying cardiac remodeling: N-Cadherin, β-Catenin and TGF-β (**C**). Counterstaining of nuclei: 4’,6-diamidino-2-phenylindole (DAPI). Images were taken at 20x magnification. Cardiac area for the respective marker in relation to number of DAPI-stained nuclei was calculated. Displayed are mean values ± SD of n=5 (sham) and n=3 (WI) animals.

### Ex vivo *tissue analysis*

After fixation (4 % paraformaldehyde for 24 h), explanted hearts were stored in PBS. *Ex vivo* diffusion tensor imaging (DTI) at 11 T was performed on four sham and three WI hearts.

For histology, 4 μm heart slices were deparaffinised, rehydrated, and heat mediated epitope retrieval (HIER, preheated target retrieval solution [Dako, #S1699], 35 min, 95 °C water bath) was performed. After cooling (in the solution, 20 min, room temperature), staining was done as published previously[7], adding 0.3 % glycine in the blocking step. Primary antibodies: Vimentin (cell signaling, #5741), desmin (abcam, #ab15200), N-Cadherin (cell signaling, #13116), β-Catenin (cell signaling, #8480), TGF-β (Thermo Fisher, #PA1-29032), CD31 (R&D Systems, #AF3628), Lyve1 (R&D Systems, # AF2125), Podocalyxin (R&D Systems, #AF1556), Prox1 (R&D Systems, #AF2727) and Reelin (R&D Systems, #AF3820). Secondary antibodies: Alexa Fluor Plus 555 (Thermo Fisher, #A31572 or #A32816). Staining with wheat germ agglutinin (WGA) Alexa Fluor™ 488 (Invitrogen, #W11261)[7] was preceded by HIER. Nuclei were counterstained (4’,6-diamidino-2-phenylindole/ DAPI [Carl Roth, #6843.1] in PBS).

Images were taken on a Zeiss Axio Scan.Z1 (Zeiss ZEN 2.6 software). Blinded analysis was performed as described previously[7].

### Statistics

Due to small sample sizes, Mann-Whitney test and Spearman correlation analysis were applied (Graph Pad Prism 8.0 [Graphpad Software]). A pathway modulation and critical component analysis was done with partial least squares modelling using the PLSM-package in R 4.3.3 and R studio 28.2.6.

## Results

### Hearts show mild myocardial structure alterations after WI

In histologic analysis of cardiac remodelling[8], cardiomyocyte diameter mirroring cardiomyocyte (CM) hypertrophy was increased, a trend towards increased levels of desmin represented cardiomyocyte remodelling, while levels of fibrosis marker vimentin were unchanged after WI (**figure 1 B**). In pathway modulation for combined data, the manifest variable (MV) CM diameter had a strong impact on the latent variable (LV) cardiac structure (**figure 3 B**).

Histologic examination of signalling pathways underlying cardiac remodelling revealed a trend towards increased N-Cadherin level (important for myofibrillar reorganization in cardiomyocytes), towards reduced β-Catenin level (involved in induction of fibrosis and cardiomyocyte hypertrophy upon increased afterload) and towards increased TGF-β level (signalling factor in cardiac fibrosis) (**figure 1 C**), with both β-Catenin and TGF-β impacting on cardiac structure for combined data (**figure 3 B**).

### Cardiac overall vessel density is increased but lymphatic vessel density is unaltered after WI

Remodelling of cardiac vessels is part of myocardial remodelling; accordingly, pan-endothelial marker CD31 level significantly increased after WI (**figure 2 A**), while levels of lymphatic vessel markers Lyve1, Podocalyxin and Prox1 were unchanged (**figure 2 B**). However, in pathway analysis, for combined data the MVs Lyve-1 and Podocalyxin levels were important for the LV ECs measured in histology (**figure 3 B**). Levels of the lymphangiocrine protein Reelin, involved in mediating recovery of heart function after MI[3], and Reelin-CD31 ratio trended towards reduction in WI, while Reelin-lymphatic marker ratios were unaltered (**figure 2 C**).

**Figure 2.**
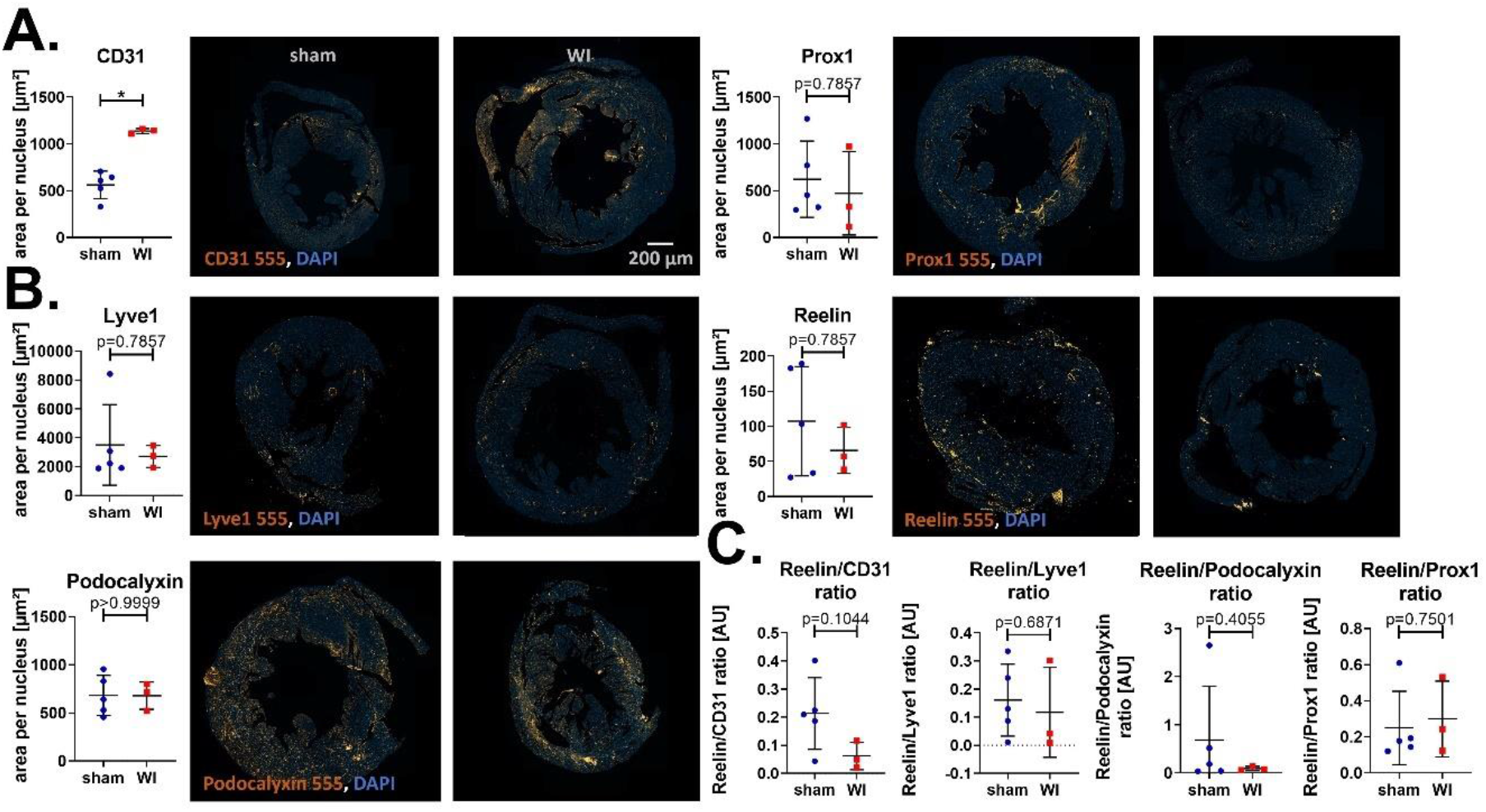
Histologic quantification of cardiac vessels: **A**) Pan-endothelial marker: CD31, **B**) lymphatic markers: Lyve1, Podocalyxin, Prox1, and **C**) lymphatic mediator Reelin. Counterstaining of nuclei: 4’,6-diamidino-2-phenylindole (DAPI). Images were taken at 20x magnification. Cardiac area for the respective marker in relation to number of DAPI-stained nuclei was calculated. Displayed are mean values ± SD of n=5 (sham) and n=3 (wire injury/ WI) animals. For statistical analysis, Mann-Whitney test was used, * = p<0.05.

**Figure 3.**
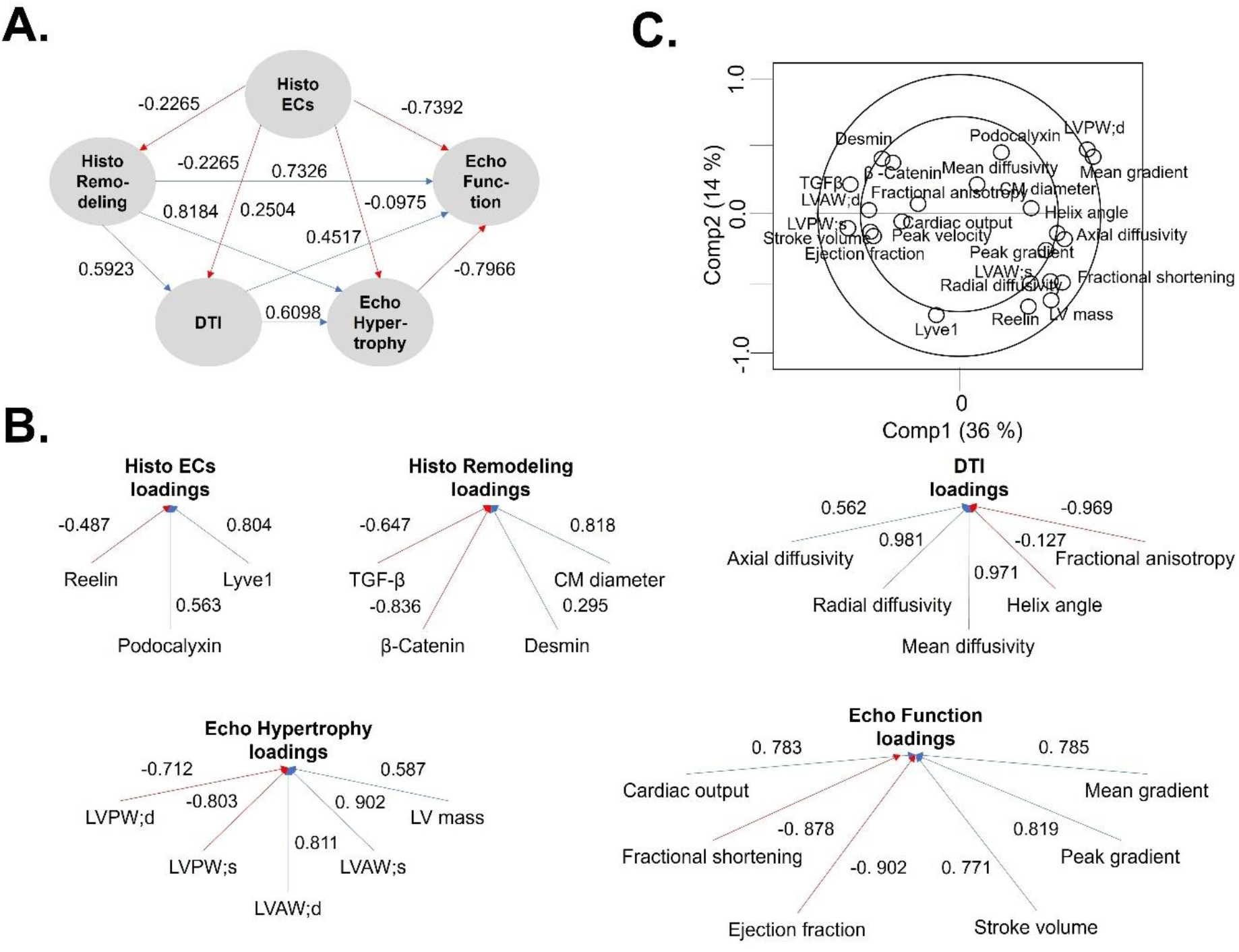
Pathway analysis with Partial least squares modelling (PLS) was performed using the PLSM-package in R 4.3.3 and R studio 28.2.6. Given are the direct effects between latent variables in the inner model (**A**) and the loading of observed variables on their respective latent variables in the outer model (**B**). **C)** Results of a critical component analysis presented as a correlation circle. DTI: diffusion tensor imaging, EC: endothelial cell, LV: left ventricle, LVAW;s/d: left ventricular anterior wall thickness in systole/diastole, LVPW;s/d: left ventricular posterior wall thickness in systole/diastole. Displayed are mean values ± SD of n=4 (sham, DTI), or n=5 (sham, histology) and n=3 (WI) animals.

### The correlation of cardiac structure parameters between DTI and histology is lost after WI

Critical component (CC) analysis of combined sham and WI groups showed a positive correlation of cardiomyocyte diameter, left ventricular mass and most DTI parameters, which were negatively correlated with desmin, TGF-β and weakly correlated peak blood flow velocity (**figure 3 C**). Reduced numbers of r-values ≤−0.5/≥+0.5, including for overall and lymphatic vessel markers for histology markers with DTI markers was found in WI, compared with sham (**figure 4 A**). Negative effects of vessel marker levels on cardiac structure measured with DTI to equal parts were direct, and indirect via cardiac structure measured in histology for combined data (**figure 3 A**). R-values for the correlation of histology markers of heart structure, and ECs were mainly positive, with an altered pattern, but unchanged average number of r-values ≤−0.5/≥+0.5 in WI compared with sham (**figure 4 A**).

**Figure 4.**
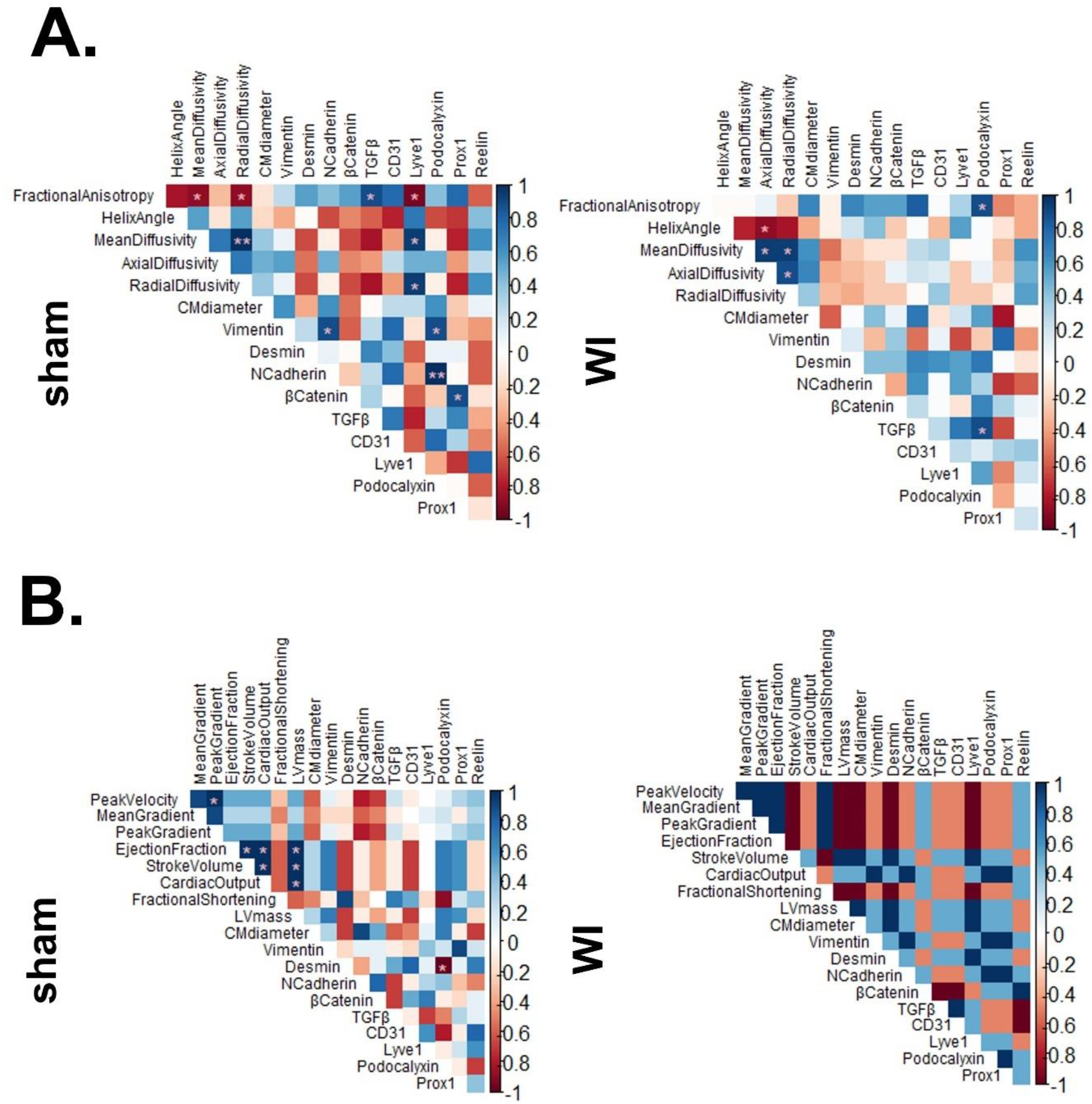
Correlation matrices for Spearman correlation of histology markers for myocardial remodeling, including vessels, with either diffusion tensor imaging (DTI) (**A**) or echocardiographic data measured at four weeks after surgery (**B**). LV: left ventricle, LVAW;s/d: left ventricular anterior wall thickness in systole/diastole, LVPW;s/d: left ventricular posterior wall thickness in systole/diastole. Displayed are mean values ± SD of n=4 (sham, DTI), or n=5 (sham, histology) and n=3 (wire injury/ WI) animals, * = p<0.05, ** = p<0.01.

### Ventricular levels of lymphatic vessel markers correlate positively with parameters of preserved heart function

Combined sham and WI data revealed a significant impact of total and lymphatic MVs on heart function parameters measured four weeks after surgery (**figure 3 A**), while CC analysis showed that Lyve-1 and Podocalyxin were indifferent towards most other variables including echocardiographic parameters. Reelin, in contrast was positively correlated with, e. g. peak gradient and LV mass, but negatively with peak velocity, stroke volume and ejection fraction (**figure 3 C**).

The correlation pattern between echocardiography and histology markers of cardiac remodeling including overall and lymphatic vessels was altered after WI (**figure 4 B**). The echocardiographic picture after WI resembled physiological cardiac hypertrophy[9]. A trend towards positive correlation of lymphatic vessel markers with cardiomyocyte diameter and left ventricular mass (signs of cardiac hypertrophy) opposed that towards negative correlation with peak blood flow velocity mirroring increased afterload as potential cause of cardiac hypertrophy. Besides, lymphatic vessel markers trended to correlate negatively with slightly reduced ejection fraction and fractional shortening mirroring mildly reduced heart function, while trending towards positive correlation with increased cardiac output and stroke volume as are found in mild cardiac hypertrophy. Except of a trend towards negative correlation with cardiac output, the pattern was similar for overall vessel marker CD31, while Reelin levels displayed opposite trends (**figure 4 B**).

Histologic structure markers directly and indirectly via DTI (**figure 3 A**), namely fractional anisotropy, mean and radial diffusivity, impacted on echocardiographic hypertrophy markers in combined data (**figure 3 B**). The LVs structural markers and ECs measured in histology impacted on heart function (**figure 3 A**).

## Discussion

This study is the first to suggest a link between ventricular lymphatic vessel density and mild loss of physiological myocardial structure and preserved heart function in a model of moderate AS. Therefore, in-depth analysis of ventricular lymphatic vasculature in future studies is important, hypothesizing that lymphatic vessels are part of early cardiac protection mechanisms upon AS. Discovery of irreversible cardiac remodelling in moderate AS, accompanied by morbidity and mortality similar to severe AS currently questions the conventional guidelines, by suggesting benefits of AVR already in moderate AS outweigh procedural risks[10], which renders knowledge about cardiac protective mechanisms crucial to allow development of alternative treatment options.

## Acknowledgements

Thanks to the team of the Heart Center Bonn for surgeries, echocardiography and kind provision of the DTI dataset.

## Ethical approval statement

Animal experiments were approved by the Landesamt für Natur-, Umwelt-und Verbraucherschutz (LANUV, Nordrhein-Westfalen, Germany) under reference number 81-02.04.2018.A250.

## Data availability statement

The data underlying this article will be shared on reasonable request to the corresponding author.

